# Automatic interpretation of otoliths using deep learning

**DOI:** 10.1101/418285

**Authors:** Endre Moen, Nils Olav Handegard, Vaneeda Allken, Ole thomas Albert, Alf Harbitz, Ketil Malde

## Abstract

The age structure of a fish population has important implications for recruitment processes and population fluctuations, and is key input to fisheries assessment models. The current method relies on manually reading age from otoliths, and the process is labor intensive and dependent on specialist expertise.

Advances in machine learning have recently brought forth methods that have been remarkably successful in a variety of settings, with potential to automate analysis that previously required manual curation. Machine learning models have previously been successfully applied to object recognition and similar image analysis tasks. Here we investigate whether deep learning models can also be used for estimating the age of otoliths from images.

We adapt a standard neural network model designed for object recognition to the task of estimating age from otolith images. The model is trained and validated on a large collection of images of Greenland halibut otoliths. We show that the model works well, and that its precision is comparable to and may even surpass that of human experts.

Automating this analysis will help to improve consistency, lower cost, and increase scale of age prediction. Similar approaches can likely be used for otoliths from other species as well as for reading fish scales. The method is therefore an important step forward for improving the age structure estimates of fish populations.

## 1. Introduction

Age of fish is a key parameter in age-structured fisheries-assessment models. Age is usually considered as a discrete parameter (age group) that identifies the individual year classes, i.e. those originating from the spawning activity in a given year (1). By definition, an individual is categorized as age group 0 from early larval stage and until January 1st, when all age groups increase the age by one. The assessment models typically express the dynamics of the individual year class from the age when they recruit, through sexual maturation, reproduction, and throughout their longevity (2). The models are fitted to data both from the commercial catches and fishery independent surveys, and a sampling program for a fish stock typically involves sampling throughout the year and across several different types of fishing gears.

The age needs to be established from the individual fish from the sampling programs. Since fish growth and, hence, age at length varies in time and space (e.g., 3) linked to environmental factors like temperature or availability of food, morphology (*e.g.,* fish length) cannot be reliably used as a proxy for age. Instead, the age is determined from a subset of individuals and usually used in conjunction with length data and information about time and location of sampling (3). The age is “read” from the annual zones in otoliths or scales. Although simple in principle, age reading depends on correct identification of zonation patterns that may consist of both true annual zones and zones representing other (unknown) temporal variation (1;4) The process is time consuming, requires a trained eye, and is uncertain. The uncertainty can be divided into accuracy and precision. Whereas the precision can be assessed by between-reader comparisons, the accuracy is difficult to identify, but may be assessed from, e.g., radiochemical analyses, analyses of chemical tags, or tag-recapture experiments (5).

Methods to automatically read otoliths have been proposed, but to date none have proven satisfactory. Fablet and Le Josse (6) investigated statistical learning techniques including neural networks and Support Vector Machines based on proposed feature extraction of images of otoliths. There are two sets of features used, biological features including fish length, sex and catch date, and geometrical features of otoliths, including shape and the opaque and translucent zonation patterns. Using both sets of features, they found that the models did not significantly improve predictions compared to using biological data only. Robertson and Alexander (7) found that precision of predicting age of otoliths using neural networks from geometric features could be improved upon by using biological features, but that the results obtained from neural networks were less precise than those obtained from experienced readers.

### 1.1. Convolutional neural network

Artificial neural networks are computational structures inspired by biological neural networks. They consist of simple computational units referred to as *neurons*, organized in layers. The parameters (or weights) of the neurons are usually learned through supervised training. This process consists of two steps, forward propagation where the network gives a prediction on the input, and back propagation where the network learns from its mistake defined by calculating the gradient of a loss function. The gradient is used to update the weights in the neurons.

In recent years, CNNs have become widely successful, especially in the field of image analysis. In 2016, the neural network designed by Krizhevsky et al. (8) was able to substantially improve performance over previous systems in the important benchmark task of object recognition, and the results were subsequently improved on by more refined network architectures (9; 10; 11) to the point of rivaling human abilities.

The most remarkable feature of deep neural networks is perhaps their *generality*. With sufficient training data, they are able to analyze and classify raw data (e.g., an array of pixels) directly. No explicit design of low-level features is necessary, the network’s lower layers learn to distinguish useful features automatically. This is achieved by structuring the network as a stack of convolutions, where each layer acts as a filter that recognizes and locates occurrences of a particular feature in the image. Subsequent layers then recognize combinations of lower layer features as more abstract features, and finally merge this information to provide a high level classification. As the same filters are applied across the whole image, an important advantage of convolutional layers is that the number of parameters to be learned is reduced, which again reduces the amount of data and computation necessary for training.

### 1.2. Objectives

Here we explore whether a convolutional neural network can be used to reliably estimate the age of an otolith from an image. We implement a network architecture and train it on otolith images from Greenland halibut (*Reinhardtius hippoglossoides*), and evaluate the precisions of our classifier by comparing it to existing age estimates from human experts.

## 2. Methods and materials

### 2.1. Data Collection

The data set consists of a set of pre-exisiting images of otoliths from the image archive of the Institute of Marine Research. Fish otoliths were collected and photographed as part of the IMR data collection program for Greenland halibut on cruises between 2006 and 2017. The otolith constitutes an important input to the stock assessment program, and represent a valuable source of historical information. There was a total of 8875 otoliths in the data set, from 4109 images of otolith pairs and 657 images of single otoliths. As the present study only investigate historical, pre-existing data and does not involve new animals, no ethics approval was deemed necessary.

The process of preparing, taking the image and reading the otoliths are described in (12). The images have a resolution of 2596 by 1944 pixels and the material contain images for both otoliths from a specimen. During preparation and transportation the otoliths are sometimes damaged or lost, and some pairs are thus not available. The images also vary somewhat in distance to object, lighting, and background. Examples of image variation and damaged otoliths are shown in Figure 1.

**Figure 1:**
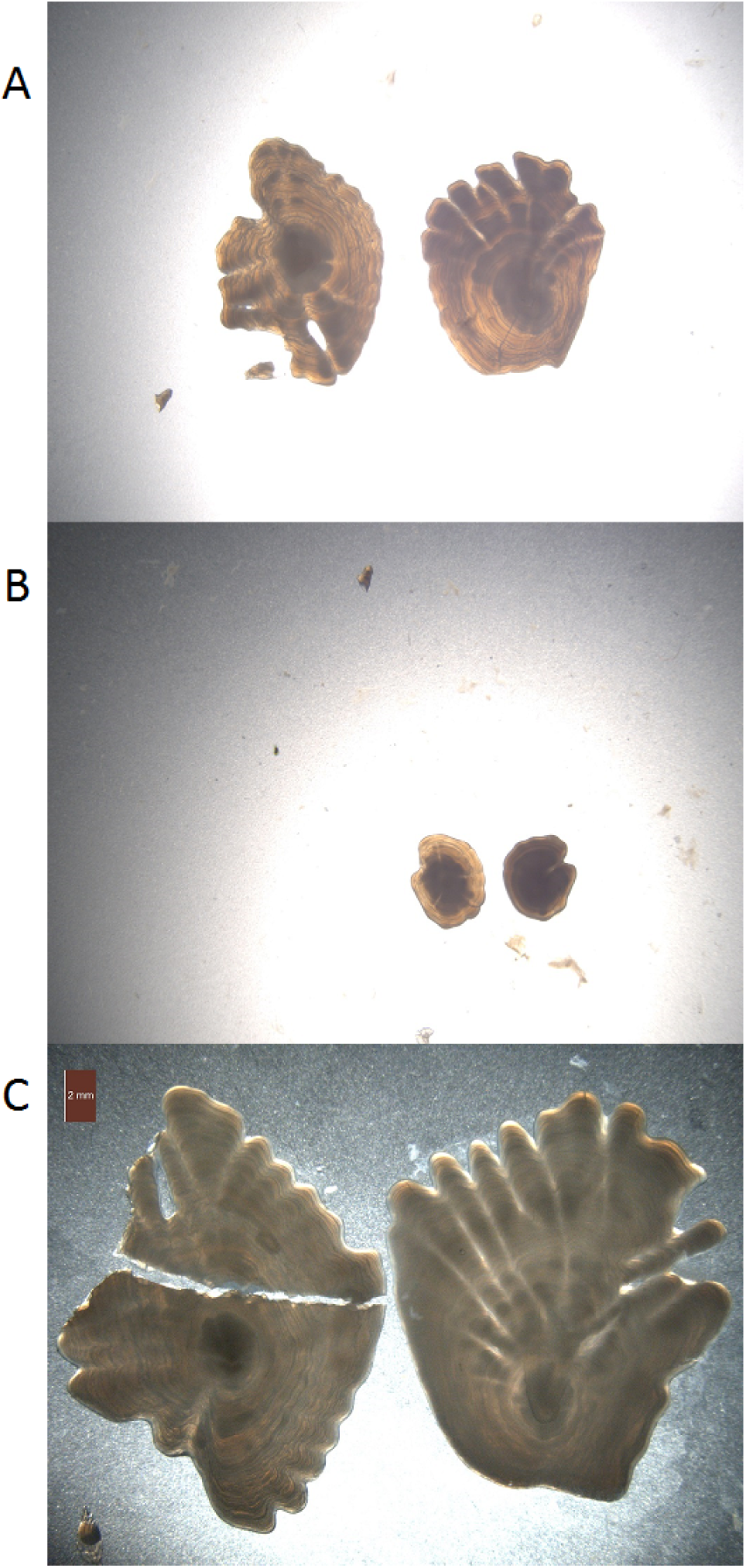
Example of otolith images. Otoliths can have loose fragments (A), vary in size (B), or are broken (C)

The age of the otoliths has been read by expert readers, and the estimated age distribution for all 8875 images is shown in Figure 2. For a long period there was no standardized method for the age reading of Greenland halibut otoliths, but two ICES workshops (13; 14) concluded with a recommendation of two different methods resulting in reasonably accurate age estimates. The age estimates here were based on the whole right otolith method (12; 14; 15). The right otolith is larger and more consistently shows pattern attributable to annuli (12). The reader is not recorded, thus there is a possibility of unknown reader biases in the data.

**Figure 2:**
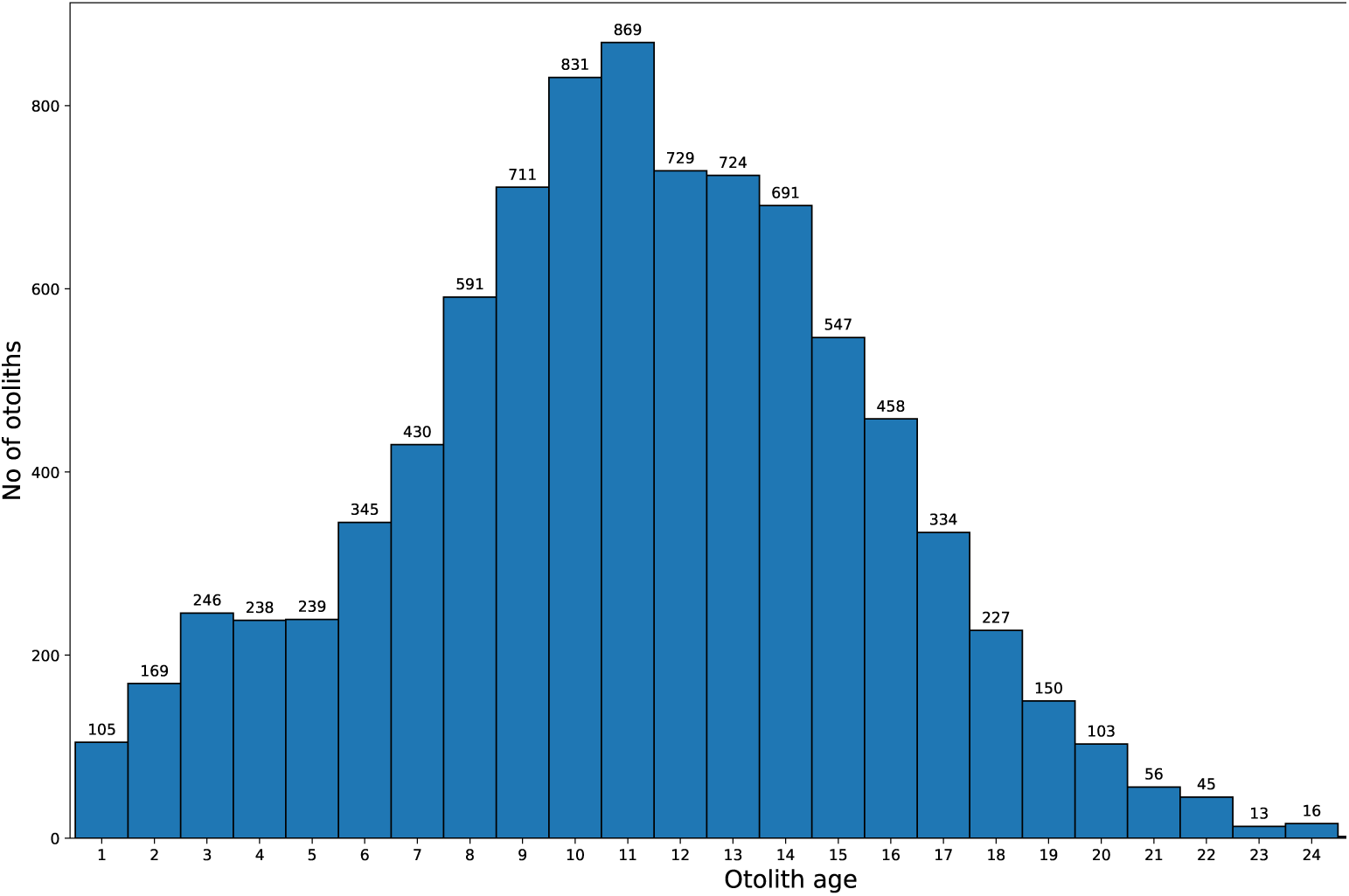
Age distribution of all 8875 images.

### 2.2. Image preprocessing

Before analysis, the images of pairs of otoliths were split, resulting in separate images of the left and right otolith. The process was complicated by variation in the placement of the otoliths, and the images were reviewed manually. The split thus varied in horizontal position up to 350 pixels. In some cases, the otoliths overlapped horizontally, and the images were allowed to overlap, resulting in images containing a small fraction of the other otolith. This overlap was rarely more than 30 pixels. Finally, images of individual otoliths were resized to a standard size of 400 by 400 pixels. Although this causes images to be stretched or shrunk, CNNs have shown to be robust to random transformations (16; 17). The process is illustrated in Figure 3. Information about image-pairs and right/left otolith was retained in order to predict the age of the pairs and evaluate the accuracy of predicting left and right otoliths.

**Figure 3:**
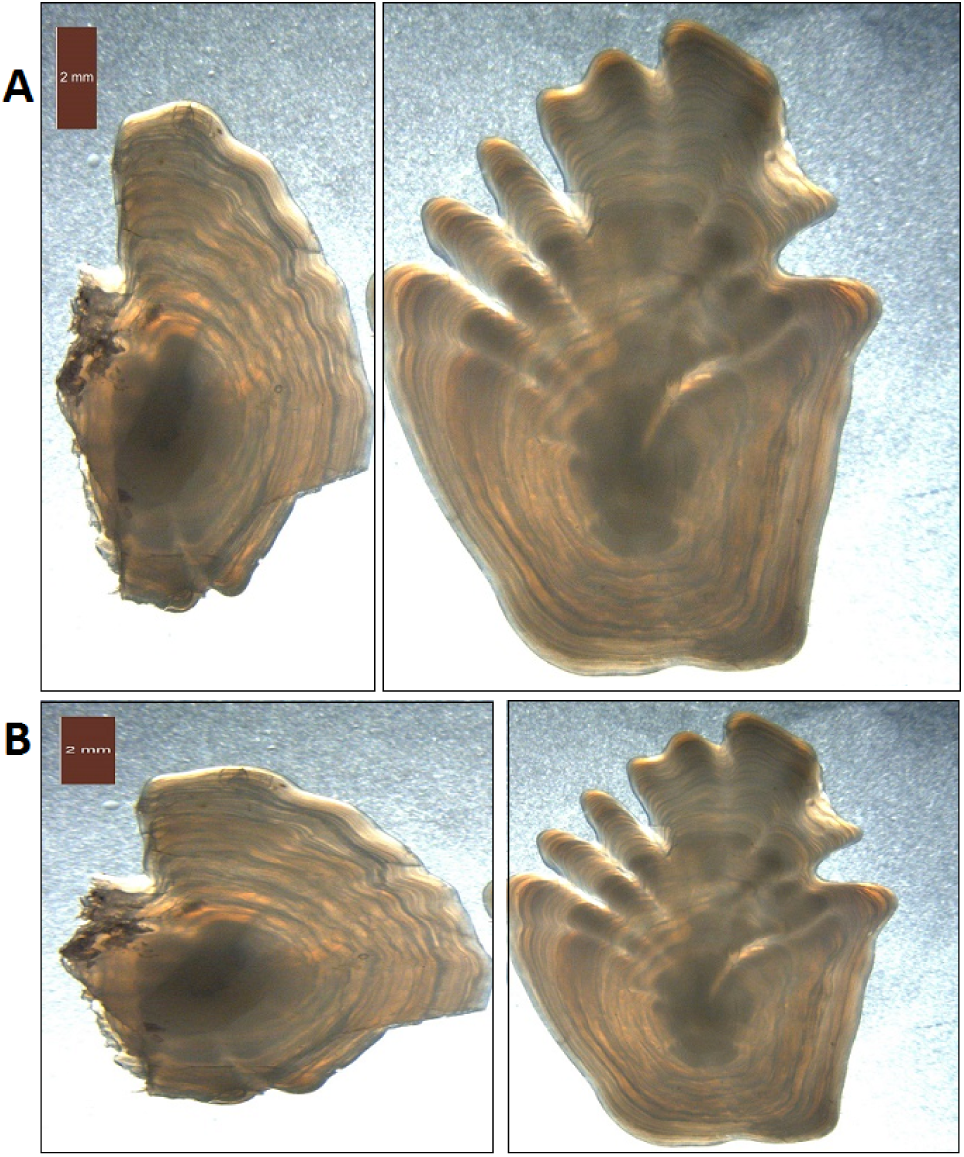
Pair of otoliths from 2014 with age estimate of 13. Due to size differences between the otoliths, the image must be split with a substantial offset from the middle. There is also a small horizontal overlap causing a fragment of the right otolith to remain in the left image. Resizing causes stretching of the images, which is particularly evident in the image of the left otolith.

### 2.3. Convolutional neural network architecture

We use a classifier model based on the Inception v3 (18) model. This is a state of the art 48-layer architecture for image classification, and the successor to the network (19) that won the 2014 ImageNet competition (20). There are several competing architectures, and variations of ResNet (9) (ResNet50, ResNet101, and ResNet152), Inception v4 (21), and DenseNet121 (22) were considered, but preliminary tests showed small differences in results, with most preliminary performance of different configurations varying less than 10%.

ImageNet classifies images of size 299×299 pixels into one of 1000 categories, and some modifications to the network was necessary. The otolith images were scaled to 400×400 pixels, and the input layer was modified accordingly. In addition, the output layer was changed from predicting classes to predicting age. Age is modeled as a regression problem, replacing the 1000-dimensional output vector with a single numeric output. The objective (or loss) function to be optimized was changed from cross entropy to mean squared error (MSE) defined as

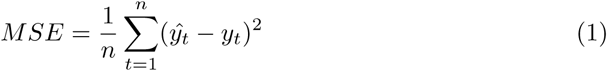

where 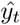 is the CNN prediction and *y*_*t*_ is the read age, and *n* is the number of predictions.

The retained layers were loaded with pre-trained, publicly available weights from training on ImageNet data. This was done to speed up training compared to random initialization (23). All layers were set to trainable.

### 2.4. Training the neural network

The network was implemented using the standard software packages Keras (24) with TensorFlow (25), and computation was performed using CUDA version 9.1 and CuDNN with nVidia P100 accelerator cards.

The data set was split into training, validation and testing sets, containing 92%, 4% and 4% of the images, respectively. The validation set is used to control (and terminate) the training process, while final accuracy is estimated using the test set. All singleton images were placed in the training set, so that the testing and validation sets only contained paired images.

Augmentation is an important technique for training deep CNNs on limited data sets (26). This process applies a set of random transformations that preserve class, artificially inflating the training data size. Thus the classifier is unlikely to see the exact same input twice, and is less likely to overfit to the data (i.e., learning to recognize individual input images, rather than identifying general features). We applied standard image augmentation supported by Keras and TensorFlow, and the images were given a random rotation between 0 and 360 degrees, reflection by the vertical or horizontal axis, and a vertically shift by +/- 10 pixels. In addition, standard image normalization for CNNs was applied resulting in mapping the 0-255 pixel values to values between 0 and 1.

The training process configuration is determined by a set of hyperparameters. Batch size defines the number of images to be processed at a time during training, and the gradient of the error function for the current parameters is calculated for each batch. The optimizer function determines how the weights are modified from the gradient. Here we used stochastic gradient descent (SGD), rmsprop, and ADAM (27). Weight decay is a regularization method that causes the weights to gravitate towards smaller values, limiting the nonlinear behavior of the classifier.

GridSearchCv from ScikitLearn (28) and KerasRegressor from Keras was used to perform a grid search of the hyperparemeter values shown in Table 1. The optimal values were found to be a batch size of 8, learning rate of 0.0004, with the ADAM optimizer (27) and a decay of 0. In addition, the patience, which controls termination of training, was set to 20, epoch was set to 150, and steps was set to 4000, so that the complete training run used 4800000 images.

**Table 1:** Hyperparameter configurations explored.

| Hyperparameter | Values explored |
| --- | --- |
| Batch size | 8, 12, 16, 20 |
| Learning rate | 0.1, 0.01, 0.0004, 0.0001 |
| Optimizer | SGD, rmsprop, ADAM |
| Weight decay | 0.01, 0.001, 0 |

### 2.5 Comparing accuracy to human experts

To compare the performance of the CNN model with that of human experts, we use the same method that are used when evaluating the humans versus humans precision (29). Since the actual age of the fish is unknown, the accuracy cannot be assessed and the Coefficient of variation (CV) between readers is used. For a given otolith *j*, reader *i* provides an age estimate *X*_*ij*_ for otolith *j*, and the CV for that individual otolith *j* is given as

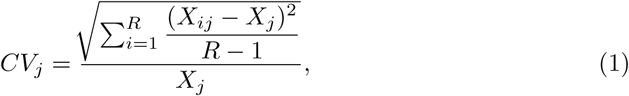

where *R* is the number of individual readers and 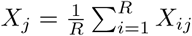. To assess the overall performance across the otoliths for the full data set, the average CV is used and defined as

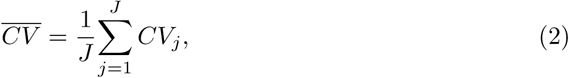

where *J* is the number of otoliths.

To evaluate the CNN model, we estimated the CV between the CNN and the human reader. Since we only have one age estimate per otolith and since we do not know the identity to the individual readers, we treated the human read otoliths as one reader and the CNN as the other. This leads to *R* = 2 where *i* = 1 is the CNN and *i* = 2 is the human reader.

Since the CNN is reading both images, we used two different definitions of the CNN read otoliths, i.e. the *X*_1*j*_ (c.f. Eq.(1)). The first uses the average over an image pair,

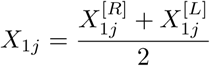

whereas the other is using the right otolith only, 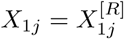where 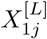and 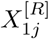is the machine predicted left and right otolith, respectively. The first is based on our approach and the latter is what an expert reader will do (12) for this specific data set, and both are tested.

## 3 Results

We conducted a series of experiments on different state-of-the-art CNNs, with varying image resolution. After prediction was made on the test set for the different configurations, the MSE of single otolith predictions was recorded. The MSE, and CV for pair-wise prediction was recorded. Then the network with the best CV was chosen.

The predictions for the left otolith (MSE = 3.25) is slightly worse than predictions overall (MSE = 3.04), and predictions for the right otoliths (MSE = 2.82) are slightly better (Table 2). However, ensemble predictions using the average prediction of the paired (left and right) otoliths (MSE = 2.78) outperform even the right otolith predictions.

**Table 2:** MSE (Eq. 1) and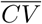 (Eq. 2) for predictions on single, left, right and paired (left and right) otolith images.

| Metric | Single | Left | Right | Paired |
| --- | --- | --- | --- | --- |
| MSE | 3.04 | 3.25 | 2.82 | 2.78 |
| $\overline{CV}$ | 9.2 | 9.8 | 8.7 | 9.0 |

From Figures 5, we see that the using both otoliths are used in an ensemble reduces prediction variance. There is also a clear tendency for the system to predict a lower age for older individuals, compared to human readers. Note that the variance between the predictions increases with the age of the otolith.

**Figure 4:**
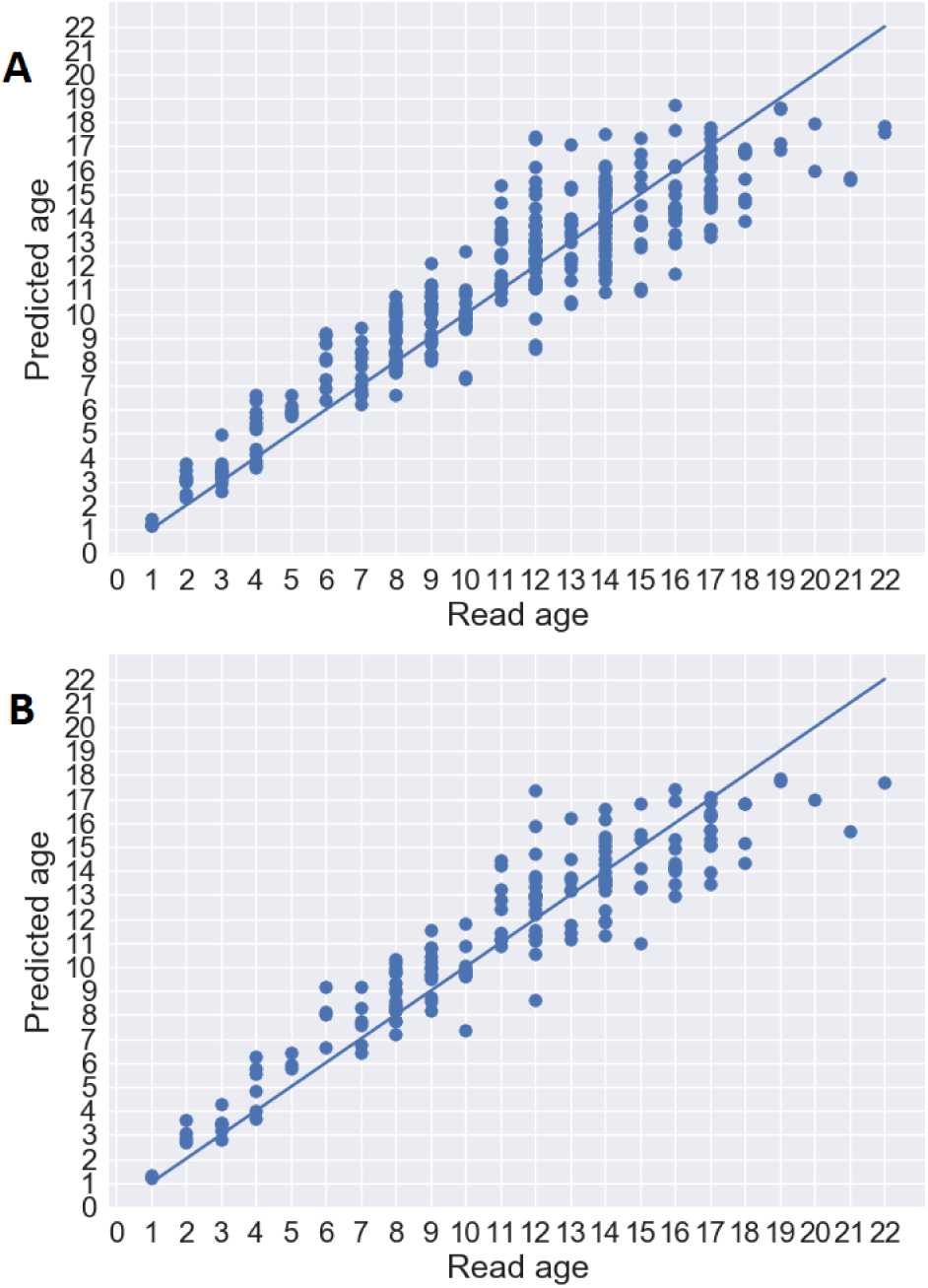
Age predictions using single otoliths (A) and when using the average prediction of each pair (B), compared to the age estimated by a human reader.

**Figure 5:**
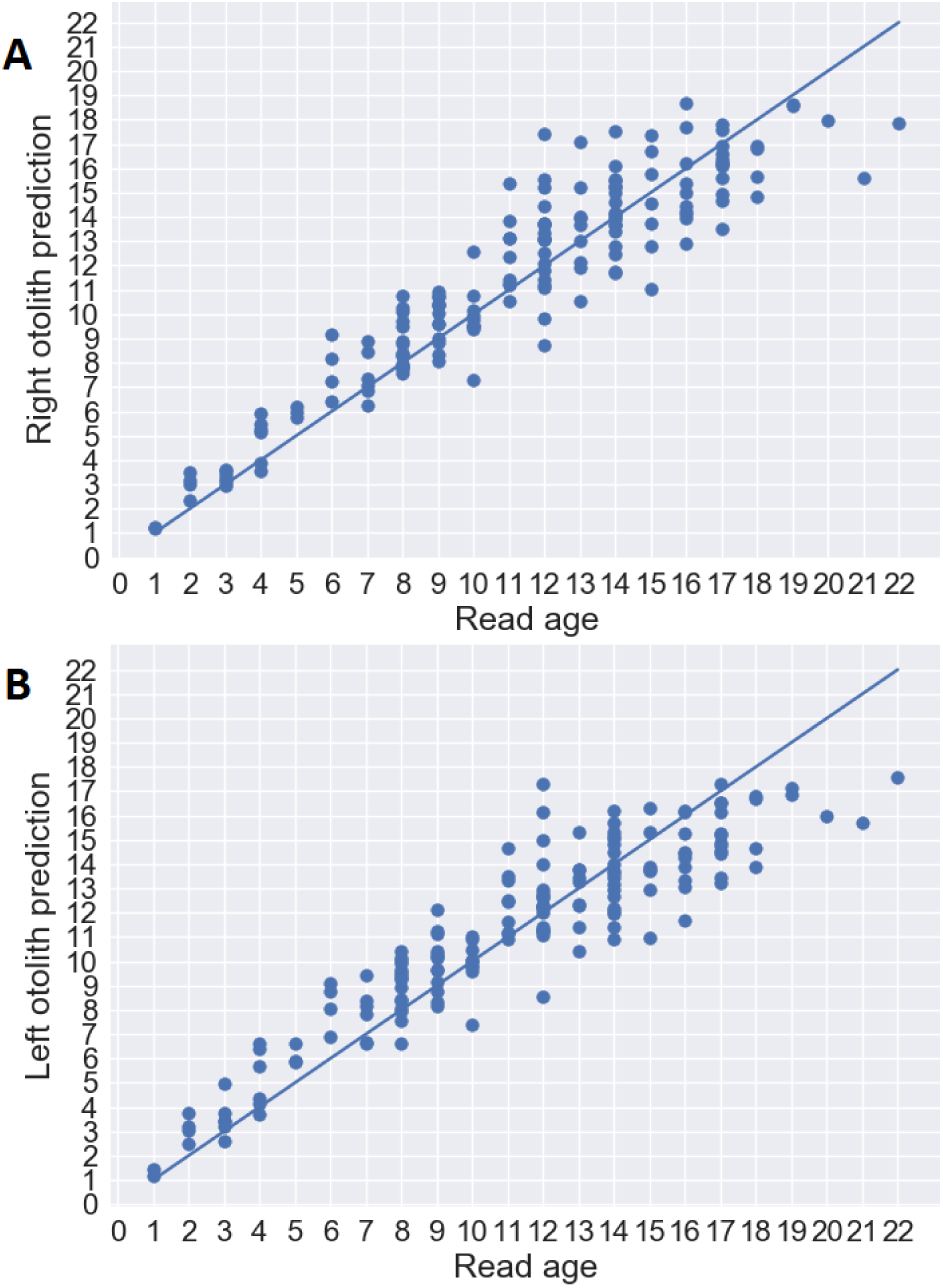
Age predictions of the right (A) and left (B) otoliths compared to the age as estimated by a human reader.

The 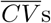 s is a commonly used measure of precision between human readers. For Greenland halibut, the 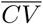 between human experts has been reported to be 12% and 16.3% (12;15). Using otolith pairs, we achieved a 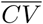 of 9.0%, while using the right otolith resulted in an 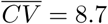 = 8.7 (Table 2). Figure 5 shows predictions for left and right otoliths separately.

## 4 Discussion

The objective of this paper was to investigate to what extent a deep convolutional neural network could be adapted to predict age from otolith images. Using a data set on Greenland halibut, we trained and validated an Inception-3 net, and showed that the network performed close to human accuracy. Deep neural networks have been shown to outperform more conventional methods on a range of problems (e.g. 26), and it is an attest to their generality that they perform well on this rather different task. Several different network architectures were tried, and most configurations were able to produce good performance, which further supports this.

The preprocessing of the images was kept as simple as possible. Potentially informative properties, such as size, proportion and orientation is lost through rescaling and augmentation, but this did not notably affect the network’s ability to predict age. The classifier also appears to be robust to varying backgrounds. Traditionally, preprocessing algorithms have also been used to improve enhance features for the classifier. We have experimented with some preprocessing techniques, e.g. we ran the images through a hill shading algorithm before training, but it did not yield better results. This supports the conventional wisdom that deep networks are able to identify informative features directly, and that efforts on developing appropriate preprocessing techniques are likely to be unnecessary.

While we have not performed an extensive analysis of the cases where the network fails to correctly predict age, but a cursory inspection revealed that image inconsistencies (some examples are shown in Figure 6) can play a part. This suggests that the results could be improved if the process of taking the images can be standardized, with consistent equipment, range, lighting, and so on.

**Figure 6:**
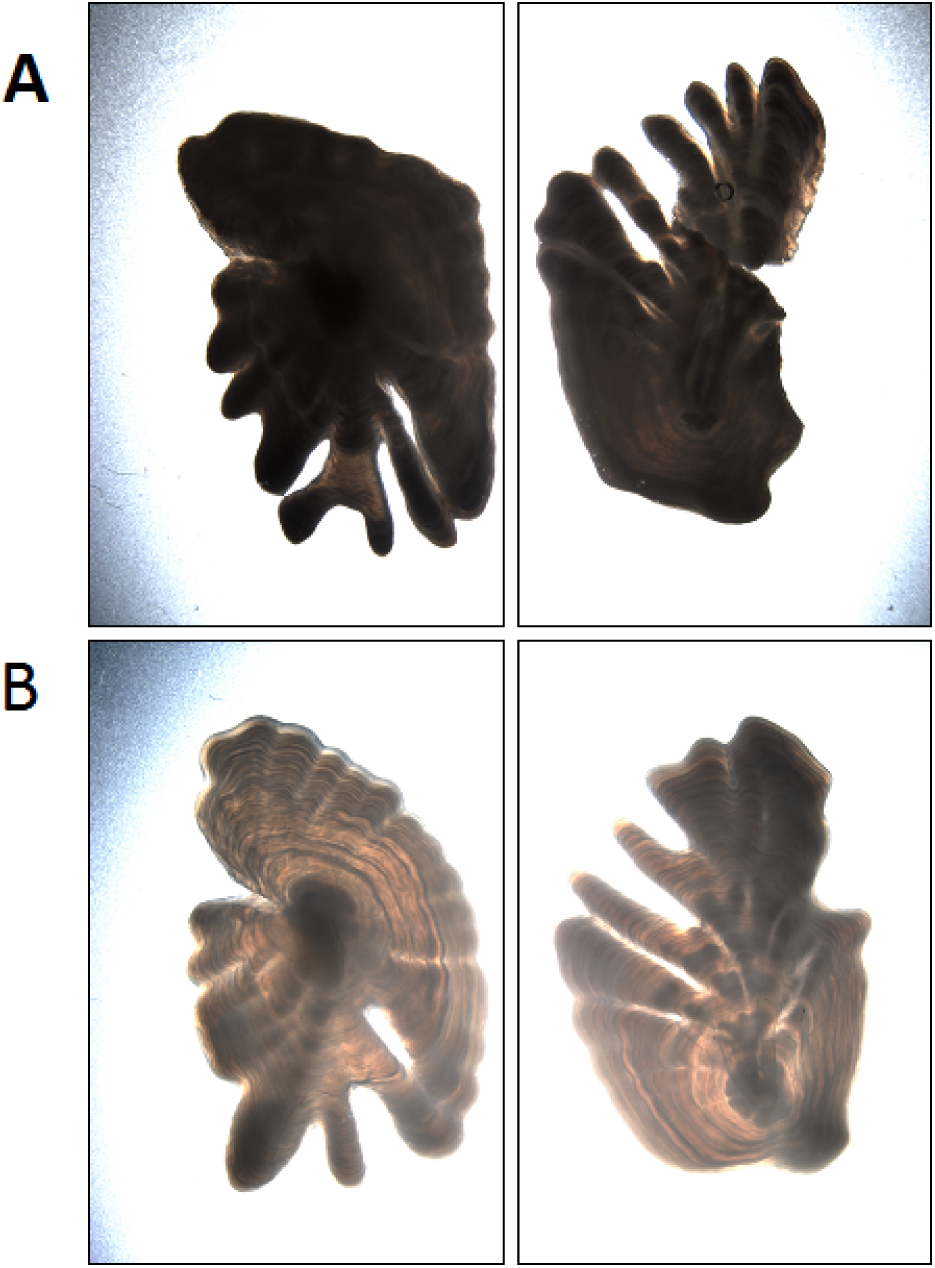
Examples of mispredicted images. A) Dark images of otoliths with deep lobes are read as 12 years, and predicted 15.7 (left), and 15.6 (right). B) Lighter otoliths below are read as 21 years, and predicted as 15.6 (left and right)

The cost function applied is not adjusted for an imbalanced data set, *i.e.,* a bias in prediction for more abundant year classes will be penalized more than for less common ones. As there are relatively few otoliths from older fish, this can be one reason for the apparent lower prediction accuracy for older otoliths. One way to mitigate this is by implementing a cost function that weight classes so that each year class inflicts the same cost (30). Such a scheme should be seen in context with actual use of the data, *e.g.*, associating higher penalties for misprediction where age determination is more critical for the assessments.

Since the model is a supervised machine learning algorithm, the learning can only be as good as the underlying precision and accuracy. Since the accuracy is unknown (12), we treated the CNN as an individual reader and computed the same mean CV as is used in human vs. human comparisons (c.f. Results). We achieved a mean CV (Eq. 2) of 9%, which to our surprise is lower than the reported mean CV between human readers of 12-16.3% (12;15). The images have been read by few readers, and it is therefore possible that individual reader bias is captured by the CNN. When using MSE as the metric, we also see that an ensemble prediction using both otoliths perform better than using the right otolith only. This could indicate that the recommended process for manual reading, which uses the right otolith only, is missing valuable information.

A common criticism to CNN’s is that we lack an understanding of the exact features used in the process. During the training and testing of the CNN we set aside 4% for both validation (during training) and testing (after training), 8% in total, that was never part of the training of the network. However, when the method is in production, it is important to keep validating the method by continuing to collect training data. This is particularly important if the method is used as part of a monitoring time-series.

Using CNNs to make age prediction can be more efficient than expert-read predictions. If cost savings are the key motivation for implementing automated aging of otoliths, a common objection is that any cost savings relies on the assumption that the actual reading is the factor that drives the cost. In reality, the time for preparing the otoliths, i.e. cutting out the otoliths and preparing the sample for imaging may take more time than the actual reading, and the saving would be marginal. However, skilled readers requires years of training, which needs to be added to the budget. Assuming that the current staff is maintained and used to generate validation data (see previous paragraph), the sampling program could be scaled up without the necessity to train more readers. Furthermore, if the network can learn features of individual readers, it is also likely that the consistency of the reading will improve, i.e. the accuracy will be lower, and the results will be reproducible, i.e. we can rerun the whole time series when the CNN is updated with new training data efficiently being able to code the expertise of a wider range of readers.

The next steps will be to test the method on new species, new features, adapt it to specific use cases and test how other predictors can be used to enhance the method. Organizing data and collecting images and age labels for a wider range of species is a key to move forward. It is likely that the patterns used to age Greenland halibut is similar to the general patterns for other species, which makes the classifier ideal for using transfer learning. One may also envision a general CNN trained on otolith images from multiple species, and then fine-tuned to each specific species. Other features like age at maturation (spawning zones) can be read from otoliths, and where training data is available, the network could be adjusted to predict these features as well.

### 4.1 Conclusion

Age determination from otoliths is an important activity for management of many marine stocks. The satisfactory result for reading Greenland halibut otoliths demonstrate that automating the data processing for this intrinsically complicated process is possible. On top of its ability to learn age-reading, the method offers improved efficiency, the possibility to (transfer) learn how to read otoliths across species, and, given proper attention to the collection of validation data, increased consistency over time. Since age is an essential component of any age based models, the method will have an impact on the management of fish resources and our understanding of ecosystem dynamics.

## 5 Acknowledgments

This project was supported by the COGMAR project, Research Council of Norway grant no 270966/O70, and by the Norwegian Ministry of Trade, Industry and Fisheries.

